# Divergent molecular signatures of regeneration and fibrosis during wound repair

**DOI:** 10.1101/2020.12.17.423181

**Authors:** Shamik Mascharak, Heather E. desJardins-Park, Michael Januszyk, Kellen Chen, Michael F. Davitt, Janos Demeter, Dominic Henn, Michelle Griffin, Clark A. Bonham, Deshka S. Foster, Nancie Mooney, Ran Cheng, Peter K. Jackson, Derrick C. Wan, Geoffrey C. Gurtner, Michael T. Longaker

## Abstract

Regeneration is the “holy grail” of tissue repair, but skin injury typically yields fibrotic, non-functional scars. Developing pro-regenerative therapies requires rigorous understanding of the molecular progression from injury to fibrosis or regeneration. Here, we report the divergent molecular events driving skin wound cells toward either scarring or regenerative fates. We profile scarring versus YAP inhibition-induced wound regeneration at the transcriptional (single-cell RNA-sequencing), protein (timsTOF proteomics), and tissue (extracellular matrix ultrastructural analysis) levels. Using cell surface barcoding, we integrate these data to reveal fibrotic and regenerative “molecular trajectories” of healing. We show that disrupting YAP mechanical signaling yields regenerative repair orchestrated by fibroblasts with activated Trps1 and Wnt signaling. Our findings serve as a multimodal map of wound regeneration and could have therapeutic implications for pathologic fibroses.

## Introduction

Fibrosis is the replacement of functional tissue with non-functional connective tissue and is the end result of damage to virtually every organ in the human body (e.g., liver cirrhosis, pulmonary fibrosis, cardiac scarring following myocardial infarction). Collectively, fibroses are implicated in up to 45% of all deaths in the U.S.(Wynn, 2004) In skin, fibrosis occurs as scarring following dermal injury, a late evolutionary adaptation to prioritize speed in wound closure. However, scars are inferior to uninjured tissue in both form and function, lacking any hair or glands and, therefore, the ability to thermoregulate or maintain skin’s normal defense/barrier function. Despite the enormous medical burden of fibrosis, no current therapies yield regenerative wound repair, due to lack of understanding of the fundamental mechanisms that would differentiate a regenerative healing pathway from a fibrotic one.(Griffin et al., 2020; Mascharak et al., 2020b; Reish and Eriksson, 2008; Walmsley et al., 2015)

We previously reported that the embryonically-derived *Engrailed-1* (*En-1*)-positive fibroblast (EPF) lineage drives dorsal dermal fibrosis in mice.(Rinkevich et al., 2015) More recently, we showed that EPFs can also arise via *postnatal En-1* activation.(Mascharak et al., 2020a) Specifically, *En-1*-negative fibroblasts (ENFs) activate *En-1* expression within the postnatal wound environment to contribute to scar tissue formation. Importantly, conversion of ENFs to pro-fibrotic EPFs is mechanically driven. Moreover, blocking mechanotransduction (via a small molecule YAP inhibitor, verteporfin) prevents this conversion and results in ENF-mediated skin regeneration, as defined by three key criteria: 1) regeneration of dermal appendages (hair follicles and glands); 2) re-establishment of extracellular matrix (ECM) architecture identical to that in unwounded skin; and 3) restoration of tensile strength to unwounded skin levels. This regenerative outcome in a typically fibrotic context posed a unique opportunity to study and define the molecular factors driving regeneration (resulting from YAP inhibition) versus fibrosis (in non-YAP-inhibited wounds).

Here, we use an integrated multi-“omics” approach to compare regenerative and fibrotic skin repair across the time course of healing at the transcriptomic, proteomic, and tissue ultrastructural levels, using single-cell RNA sequencing (scRNA-seq); timsTOF proteomics (a high-resolution shotgun sequencing technology able to identify over one thousand unique peptides from a small sample); and a unique algorithm for quantifying ECM ultrastructure, respectively. We employ a cell-barcoding system to correlate data across modalities at single-mouse resolution, allowing us to impute bulk (proteomic, ECM ultrastructure) data onto our single-cell (scRNA-seq) dataset and to achieve significantly increased depth of integration across all three high-dimensional datasets in pseudotime. We reveal two repair trajectories defined by distinct molecular motifs in fibroblasts: one “fibrotic,” dominated by mechanical signaling motifs; and one “regenerative,” characterized by activation of stem cell- and development-related pathways. By elucidating the divergent signatures of regeneration and fibrosis at the transcriptomic, proteomic, and tissue ultrastructural levels across all key phases of tissue repair, we identify the biology underlying a novel example of adult mammalian wound regeneration. Our work represents a multi-“omic” roadmap of tissue repair, which can serve as a resource for future studies into regeneration and fibrosis.

## Results

The injury response is defined by multiple phases with distinct cell types and activities (**Figure S1**); it is therefore critical to examine the cellular and molecular dynamics of healing tissue over time. We thus analyzed wounds and skin at five separate timepoints: unwounded skin; postoperative day (POD) 2, when inflammatory processes dominate; POD 7, the peak of fibroblast proliferation; POD 14, when wounds are fully re-epithelialized and fibroblasts are actively producing new ECM; and POD 30, when fibroblasts are engaged in ECM remodeling. To study the dynamics of tissue repair in a clinically relevant fashion, we used a splinted excisional mouse wounding model, which prevents the rapid wound contraction typical of loose-skinned mice, instead inducing human-like healing kinetics and formation of a fibrotic scar.(Galiano et al., 2004) Immediately following wounding, the wound bed was injected with either verteporfin or vehicle-only (PBS) control. Consistent with the typical postnatal healing outcome, control wounds formed distinct fibrotic regions of scar, which grossly appeared as pale, bare (hairless) areas, and on histology exhibited dense, linearly-aligned ECM fibers without secondary elements such as hair follicles or glands (**Figure S1**, left). In contrast, verteporfin-treated wounds healed via regeneration of unwounded skin structure, with gross hair regrowth, glands, and less-dense, more randomly-oriented ECM fibers (**Figure S1**, right).

In order to compare the cellular and molecular dynamics of wound repair in verteporfin-treated versus control wounds, five mice per timepoint (unwounded and POD 2, 7, 14, and 30) and treatment condition (control and verteporfin) were wounded (2 wounds pooled per animal) and analyzed. For each individual mouse, wound tissue was divided into thirds. One-third of the total tissue from each mouse was processed for histology (see **Figure 1A**); the remaining tissue was processed for proteomic and scRNA-seq analysis. Tissue was digested to obtain cells, which were sorted using an established FACS strategy in order to isolate fibroblasts (Lin-) and all other cells (primarily immune cells; Lin+). Half of the FACS-sorted cells were subjected to proteomic analysis; peptides were purified separately from Lin- and Lin+ cells from each mouse and subjected to timsTOF mass spectrometry. The remaining half of FACS-sorted cells were subjected to scRNA-seq analysis; we tagged each sample using a hashtag oligonucleotide cell surface barcoding system, with unique 12-base oligomers used to track the individual mouse and cell type (Lin-vs. Lin+) for each sequenced cell. Barcoded cells from up to 10 samples were pooled per sequencing run, then captured for scRNA-seq analysis using the 10x Chromium platform. Data demultiplexing was employed to identify the mouse of origin for each single cell transcriptome, enabling linkage of scRNA-seq samples with companion proteomic and histologic samples derived from the same mouse.

**Figure 1.**
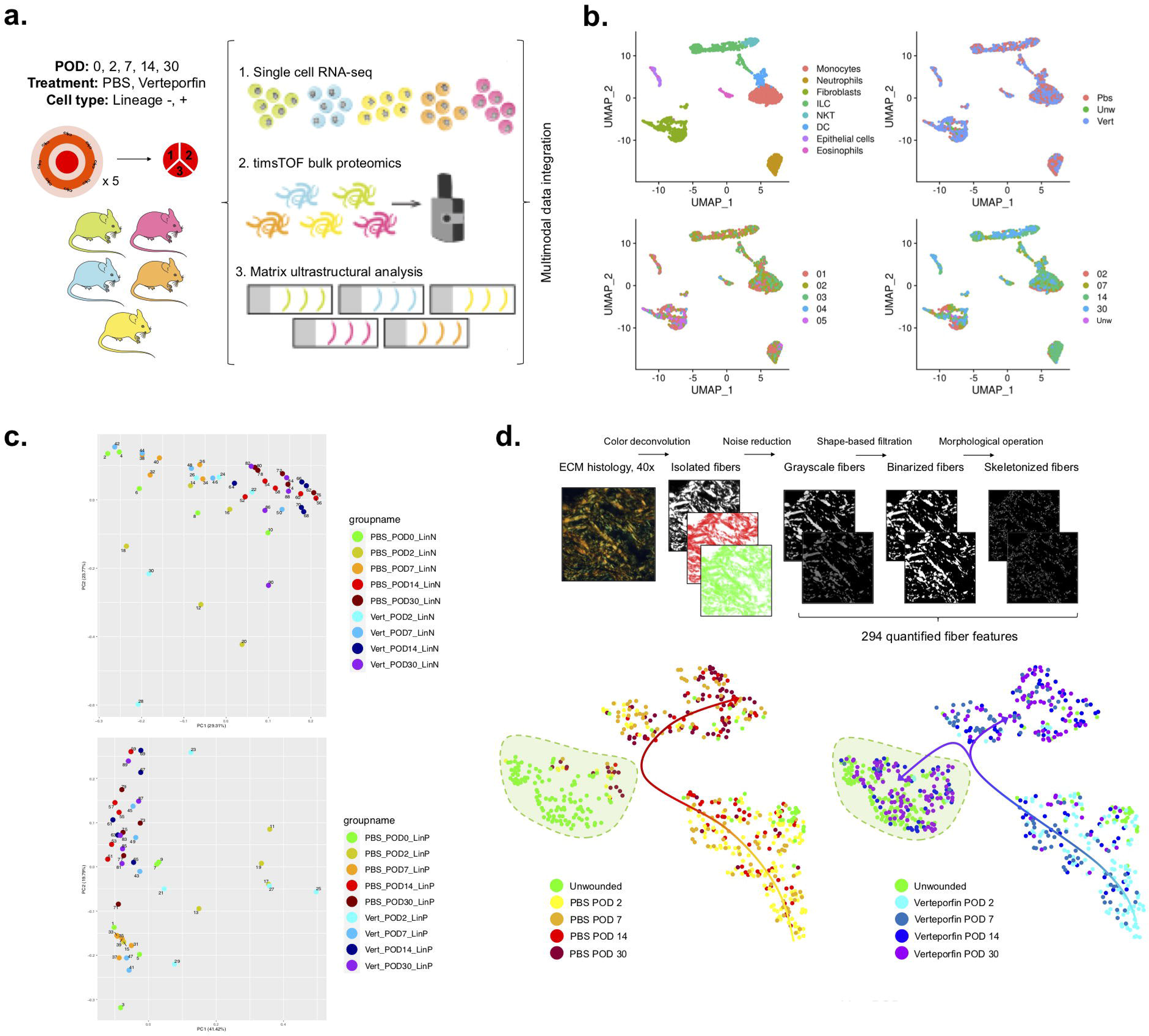
Multimodal interrogation of scarring and regenerative skin wound repair. **(A)** Experimental strategy for mouse-paired single-cell RNA-sequencing (scRNA-seq), timsTOF bulk proteomics, and matrix ultrastructural analysis of wound healing. **(B)** Manifold projection of scRNA-seq cells colored by cell type (top left), treatment group (top right), individual mouse (bottom left), and postoperative day (POD, bottom right). **(C)** Principal components mapping of bulk timsTOF proteomic data for Lin-(fibroblasts, top) and Lin+ (non-fibroblast, bottom) cells. **(D)** Top: Schematic outline of image processing and analysis pipeline to quantify 294 matrix parameters. Bottom: t-SNE plots of control (left) and verteporfin (right) wounds at POD 2, 7, 14, and 30. Green shaded region in both plots corresponds to unwounded skin cluster. Overlaid arrows reflect evolution of ECM over time.

For scRNA-seq, across all timepoints, treatment conditions, and cell types, we analyzed a total of 2,986 cells (**Figure 1B**). Cells formed distinct transcriptomic clusters in UMAP-space based on cell identity (**Figure 1B**, top left); using data demultiplexing, we also analyzed transcriptomic clusters based on mouse of origin (**Figure 1B**, bottom left), treatment condition (**Figure 1B**, top right), and postoperative timepoint (**Figure 1B**, bottom right). Cell proportions were similar between control- and verteporfin-treated wounds at all timepoints, suggesting that verteporfin may induce regeneration by fundamentally altering cell phenotypes rather than relative representation of different cell types, or alternatively may alter representation of subpopulations of these canonical cell types (**Figure S2A**). To address if selective changes occurred at the protein level, we next examined timsTOF proteomic profiles of Lin- and Lin+ cells from wounds treated with either verteporfin or PBS, with a focus on highly enriched markers reflecting changes in fibrotic state. We observed notable increases in Col2a1 and Col6a2 by POD 30, as well as significant changes in other fibrotic markers such as Col1a1, Diablo, and Cdh11 that selectively changed in response to verteporfin treatment. Using principal component analyses, we found that Lin- and Lin+ proteomic specimens differed strongly by time point (**Figure 1C**, top and bottom, respectively), reflecting key changes in the tissue repair process (many linked to cytoskeletal reorganization). Overall, differences between PBS- and verteporfin-treated specimens were more narrowly focused (**Figure S3A and S3B**), suggesting the proteomic analysis could capture coherent fibrotic changes among a broader program of dynamic reparative signals.

Finally, we quantified ECM ultrastructure from skin and wounds using a newly developed image processing algorithm (**Figure 1D**, top). Matrix architecture is a critical determinant of tissue’s physical properties; however, differences in ECM structure/organization may not be directly evidenced by differences in transcription or translation of ECM component proteins, making such quantitative characterization of ECM ultrastructure (and correlation of these tissue-level features to upstream transcriptomic/proteomic features) an important aspect of analysis. Using this computational pipeline, we quantified 294 parameters from connective tissue histology sections throughout each wound, then performed clustering analysis via t-SNE to compare samples based on these parameters. ECM from control wounds across time diverged from, and exhibited almost no overlap with, unwounded skin ECM (**Figure 1D**, bottom left), demonstrating that typical wound healing results in ECM ultrastructure fundamentally distinct from the “normal” unwounded skin architecture. In contrast, verteporfin-treated wounds at later timepoints (POD 14 and 30) largely overlapped with the unwounded skin cluster (**Figure 1D**, bottom right), suggesting that healing following verteporfin treatment resulted in ECM ultrastructure similar to that of unwounded skin. In both treatment groups, the most significant differences in ECM structure were found between POD 7 and 14, consistent with ECM synthesis and deposition largely occurring between these timepoints.

While all three data modalities demonstrated clustering primarily by differences in biological conditions (timepoint, treatment), they also showed heterogeneity between individual mice. Transcriptomic signatures were generally most variable between mice in unwounded skin and POD 2 and became increasingly homogeneous at later wound timepoints (**Figure S2B and S2C**). Bulk proteomic data were broadly more heterogeneous, with considerable variability between mice at POD 2 and 7 (**Figure 1C, Figure S3C and S3D**). In contrast, matrix ultrastructural data were most heterogeneous at POD 7 and 14, during the stages of active matrix deposition (**Figure 1D, Figure S4B**).

Having established our ability to profile wound dynamics using single-cell transcriptomic, proteomic, and ECM ultrastructural analysis, we next sought to apply these three methods to identify divergent mechanisms of regeneration versus fibrosis throughout wound repair. We analyzed three cell populations relevant to wound repair: fibroblasts; myeloid lineage cells (granulocytes, monocytes, macrophages); and lymphocytes (NK cells, T cells, B cells). As fibroblasts are highly mechanosensitive cells as well as the primary cellular drivers of fibrosis,(Aarabi et al., 2007; Barnes et al., 2018; Wong et al., 2011) we hypothesized that verteporfin primarily promotes regeneration by altering fibroblast phenotype. We thus focused our analysis on the transcriptionally-defined fibroblast subgroup from our scRNA-seq data (**Figure 2A**). We used Monocle3 to generate temporally-informed but *treatment-agnostic* trajectories among cells within our dataset. Based on inspection, an optimal manifold was selected that maximally captured changes between postoperative days; importantly, inspection and choice of manifolds was performed in an entirely treatment-agnostic fashion to minimize bias. The resulting trajectory was characterized by two bifurcating branches, with early-timepoint (POD 2) wound fibroblasts clustered near the branchpoint and late wound fibroblasts found at either end of the two branches (**Figure 2B**, top panel). Interestingly, unwounded skin fibroblasts were almost exclusively found in the bottom branch (**Figure 2B**, second panel), which was also relatively enriched with fibroblasts from verteporfin-treated wounds. Given these findings, we postulated that this bottom arm represented a comparatively regenerative wound repair trajectory, with fibroblasts progressing toward an unwounded skin-like transcriptional state, while the upper arm (enriched in PBS-treated wound fibroblasts and lacking unwounded skin fibroblasts) represented a more fibrotic trajectory. Of note, while the regenerative and fibrotic arms were enriched for wound cells from verteporfin-treated and control wounds, respectively, some control wound fibroblasts were also found within the regenerative arm (**Figure 2B, second panel**). This finding highlights the possibility that wound regeneration and fibrosis are not, at the cellular level, mutually exclusive processes; rather, cells with pro-regenerative activity may be present within even scarring wounds, but in the absence of intervention (e.g., YAP inhibition), the mechanically-activated, pro-fibrotic molecular programming may override and lead to the dominant scarring phenotype.

**Figure 2.**
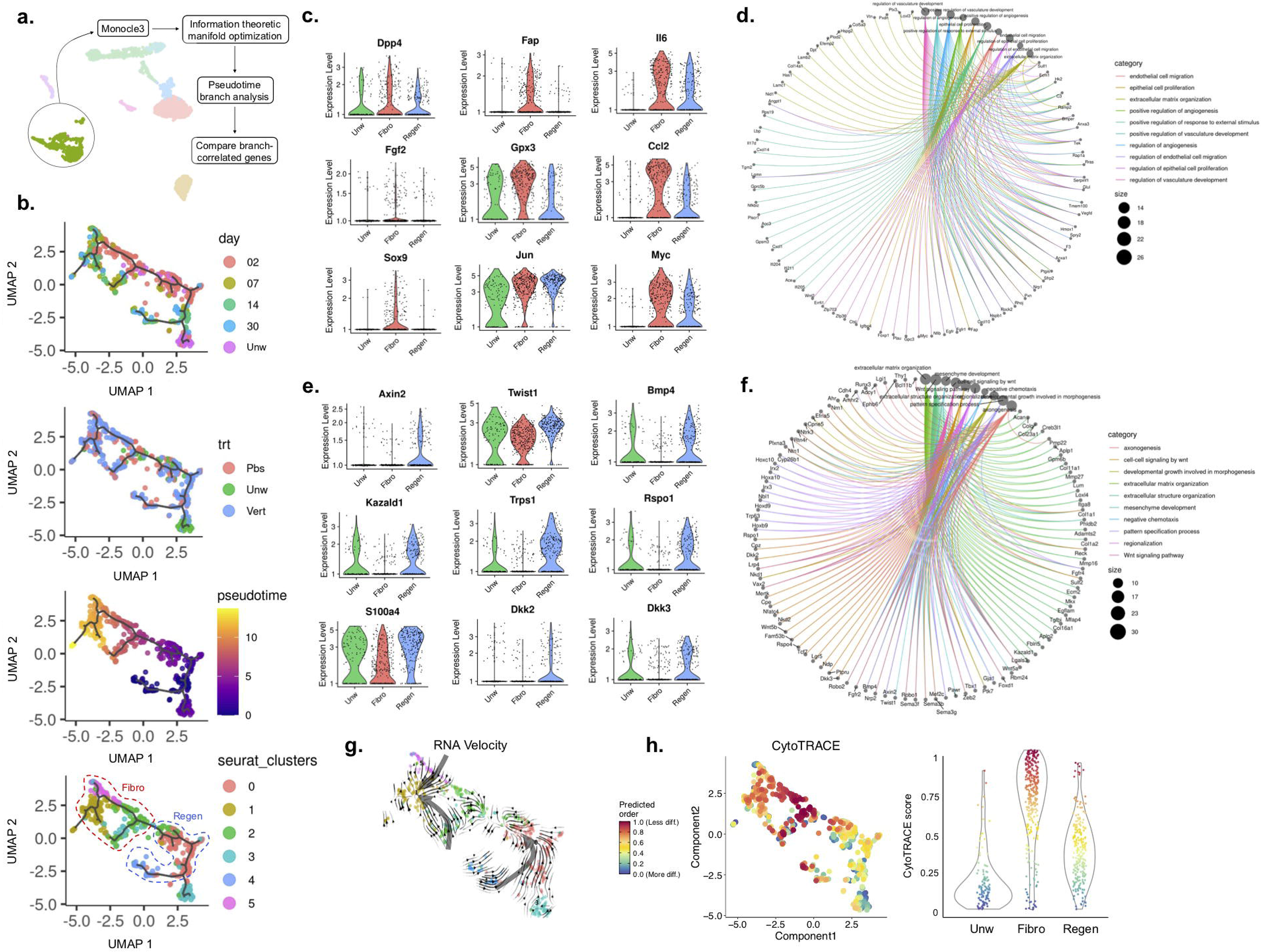
Transcriptomic trajectory analysis of fibroblasts in pseudotime. **(A)** Schematic outlining manifold generation, optimization, and branchpoint analysis for fibroblasts in Monocle3. **(B)** Fibroblast manifold colored by POD (top), treatment group (upper middle), pseudotime value (lower middle), and Louvain-based (Seurat) cluster (bottom). **(C,D)** mRNA expression of selected genes (C) and functional enrichment analysis of genes (D) positively correlated with pseudotime. **(E,F)** mRNA expression of selected genes (E) and functional enrichment analysis of genes (F) negatively correlated with pseudotime. **(G)** RNA velocity analysis of fibroblasts. **(H)** CytoTRACE analysis of fibroblasts, with manifold projection colored by CytoTRACE score (left) and CytoTRACE scores for cells in each arm of the manifold (right).

When we performed similar analysis on myeloid and lymphoid cells, none of the manifolds generated exhibited similar bifurcation or divergence based on treatment condition (**Figures S5 and S6**). Pseudotime trajectory analysis of myeloid cells (**Figure S5A-C**) demonstrated enrichment for pro-inflammatory genes (e.g., *Ccl6*, *Ccl9*, *Cxcl2*) at early timepoints and enrichment for multiple genes involved in antigen presentation (e.g., H2 complex proteins) at later timepoints (**Figure S5C-E**). However, these genes were similarly expressed in PBS and verteporfin wound cells, indicating that they did not differentiate fibrotic versus regenerative healing outcomes. Similarly, for lymphoid cells, pseudotime analysis (**Figure S6A-C**) did not significantly differentiate between fibrotic and regenerating wound cells and instead reflected early (**Figure S6C and S6E**) and late (**Figure S6C and S6D**) infiltration of natural killer (NK) and B/T lymphocytes, respectively. Collectively, these data were consistent with verteporfin primarily exerting its pro-regenerative effects by acting on fibroblasts, rather than modulating the immune response.

Given the identification of putative fibrotic and regenerative cell trajectories in the fibroblast manifold, we sought to identify distinct signatures of these two fibroblast trajectories. Louvain-based (Seurat) clustering of all fibroblast cells identified a total of six clusters: clusters 0 and 4 collectively made up the “regenerative” arm; clusters 1, 2, and 5 made up the “fibrotic” arm; and cluster 3 primarily comprised unwounded skin fibroblasts (**Figure 2B**, bottom panel). We then calculated pseudotemporal Pearson correlations for each gene along our primary trajectory of interest (**Figure S7A**). We found that the fibrotic arm and its clusters (1, 2, and 5) were enriched for expression of known “pro-fibrotic”/“activated” fibroblast markers such as *Dpp4* (CD26, which marks the scarring *En1-*positive fibroblast lineage),(Rinkevich et al., 2015) *Fap*, and *Gpx3* (**Figure 2C, Figure S7B**). Fibrotic trajectory cells also had increased expression of *Fgf2*, *Il6*, *Ccl2, Sox9*, and *Myc,* whose upregulation in fibroblasts has been implicated in focal adhesion kinase (FAK)-mediated organ fibrosis and tumor stroma deposition.(Murphy et al., 2020; Wong et al., 2011) Gene set enrichment analysis on the top 1% of genes positively correlated with pseudotime revealed terms related to ECM deposition (e.g., ECM structural constituent, GAG binding), growth factor production/activity, and mechanotransduction signaling (e.g., cell adhesion molecule binding, integrin binding), as well as fibrosis-related Reactome terms such as MAPK signaling, Ras signaling, and focal adhesion (**Figure 2D, Figure S7D**), supporting a classically activated/pro-fibrotic phenotype characteristic of *Engrailed-1* lineage-positive fibroblasts.

We next examined the genes most strongly anticorrelated with this pseudotime trajectory. Interestingly, consistent with the more regenerative phenotype whose cells were over-represented along this branch, we identified multiple genes involved in developmental and regenerative pathways (**Figure 2E, Figure S7C**). These included genes previously implicated in regenerative tissue response to injury, including *Bmp4* (known to regulate mammalian digit tip regeneration)(Han et al., 2003) and *Kazald1* (implicated in axolotl limb regeneration and blastema formation).(Bryant et al., 2017) We also identified several genes involved in Wnt signaling (known to be important in hair follicle growth and development),(Ito et al., 2007) including Wnt signaling targets *Axin2*, *Twist1,* and *S100a4*; Wnt pathway agonist *Rspo1*; *Dkk2* and *Dkk3*, key factors in embryonic skin development;(Song et al., 2018) and Wnt signaling pathway regulator *Trps1.* The latter was of particular interest as *Trps1* has been identified as a putative “master regulator” of Wnt signaling and is specifically implicated in hair follicle morphogenesis and proliferation as well as human hypertrichotic syndromes (characterized by excessive hair growth).(Fantauzzo and Christiano, 2012; Fantauzzo et al., 2012; Fantauzzo et al., 2008) By gene set enrichment analysis on the top 1% of genes anticorrelated with pseudotime, the regenerative arm was enriched for BMP and Wnt signaling-related terms such as Frizzled binding (**Figure 2F, Figure S7E**). Enriched Reactome terms also included multiple Wnt signaling-related terms, as well as “signaling pathways regulating pluripotency of stem cells.” These unique functional enrichment terms, combined with the observation that fibroblasts in this trajectory expressed lower levels of *Dpp4* (which marks EPFs; **Figure 2C,**top left panel), implied a molecularly distinct trajectory of healing orchestrated by non-scarring, *Engrailed-1* lineage-negative fibroblasts.

We sought to further interrogate the differences underlying these putative trajectories using RNA velocity analysis. This approach leverages the principle that higher representation of unspliced mRNA variants is correlated with increased “speed” of mRNA transcription.(Bergen et al., 2020) Based on relative prevalence of spliced and unspliced mRNA variants, future transcriptomic states of individual cells can be predicted in a directional manner and in relation to other cells within the same dataset, enabling inference of molecular dynamics among cell populations. Applying this analysis revealed that the fibrotic and regenerative arms had opposing RNA velocity directionalities; overall, RNA velocity vectors in the fibrotic arm pointed in the direction of increasing postoperative time, while those in the regenerative arm were directed *away* from later postoperative days and *toward* cells from earlier healing timepoints (**Figure 2G**). We also performed analysis with CytoTRACE, a computational pipeline that predicts individual cells’ relative differentiation states from scRNA-seq data based on the distribution of unique mRNA transcripts.(Gulati et al., 2020) CytoTRACE analysis revealed that unwounded skin fibroblasts were the most differentiated (corresponding to lower CytoTRACE scores), while healing wound fibroblasts were relatively less differentiated (corresponding to higher CytoTRACE scores, Figure 2H left panel). Furthermore, fibroblasts in the fibrotic trajectory were less differentiated than those in the regenerative trajectory (**Figure 2H** right panel). Within the fibrotic trajectory, progressing from earlier to later healing timepoints, fibroblasts exhibited an overall increase in degree of differentiation (decreasing CytoTRACE scores; **Figure 2H**and **Figure S7F** top row). In contrast, cells in the regenerative branch appeared to *decrease* in degree of differentiation along their trajectory, suggesting increased developmental potential as these cells progressed through the wound repair process (**Figure 2H** and **Figure S7F** bottom row).

In order to further elucidate changes in cell states along each trajectory, we performed CytoTRACE analysis independently on the fibroblast clusters comprising each trajectory and examined the genes most significantly correlated with differences in CytoTRACE scores along each branch. Decreasing CytoTRACE scores across time in the fibrotic trajectory were largely driven by ECM-related genes (**Figure S7F** top row), consistent with differentiation *toward* a mature scar fibroblast fate; conversely, increasing CytoTRACE scores across time in the regenerative arm were explained by de-differentiation *away* from a scar fibroblast fate (**Figure S7F** bottom row). Interestingly, in verteporfin-treated (regenerative) wounds, features of regeneration such as hair follicle regrowth and restoration of normal ECM ultrastructure were most prominent in later stages of wound healing (e.g., POD 14 and 30), consistent with progression toward a less-differentiated and increasingly developmental-like fate demonstrated by CytoTRACE. Collectively, CytoTRACE findings supported our observation of opposing trajectories of cell molecular dynamics based on RNA velocity analysis, and independently confirmed the distinct fibroblast phenotypes in the fibrotic versus regenerative arm.

For our cross-platform analysis, we first broadly examined our proteomic data from fibroblasts (Lin-cells). Leveraging the aforementioned mouse-to-mouse variability (**Figure S3**), we used imputation analysis to map protein expression values onto cells in our scRNA-seq dataset based on shared mouse of origin. Consistent with the transcriptomic profile of the fibrotic arm, we found that expression levels of proteins involved in mechanical signaling, such as Ank1 (a well-validated readout of YAP activity)(Elster et al., 2018) and Dag1, were also higher in the fibrotic trajectory at POD 2 and 7 (**Figure 3A**). Gene set enrichment analysis of the proteins with expression most strongly enriched in the fibrotic arm revealed terms related to growth factor signaling, mechanical signaling including integrin binding and cell adhesion, and contractility-related processes (**Figure 3B**, top panel). In addition, ECM deposition and organization were strongly enriched, including multiple ECM component proteins such as collagens (Col1a1/2, Col2a1, Col3a1, Col6a1), fibulin (Fbln1), and fibronectin (Fn1). These integrated proteomic findings supported the highly mechanically activated and strongly ECM-producing scar fibroblast phenotype demonstrated by our scRNA-seq data. In contrast, gene set enrichment analysis of proteins most strongly correlated with the regenerative arm revealed a lack of mechanical signaling-, contractility-, or ECM-related terms, consistent with scRNA-seq data for this arm (**Figure 3C and 3B bottom panel**). Overall, the regenerative arm was significantly less enriched in contractile and ECM-producing proteomic signatures than the fibrotic arm, consistent with the lack of a mechanically-activated fibroblast transcriptomic signature in the regenerative arm.

**Figure 3.**
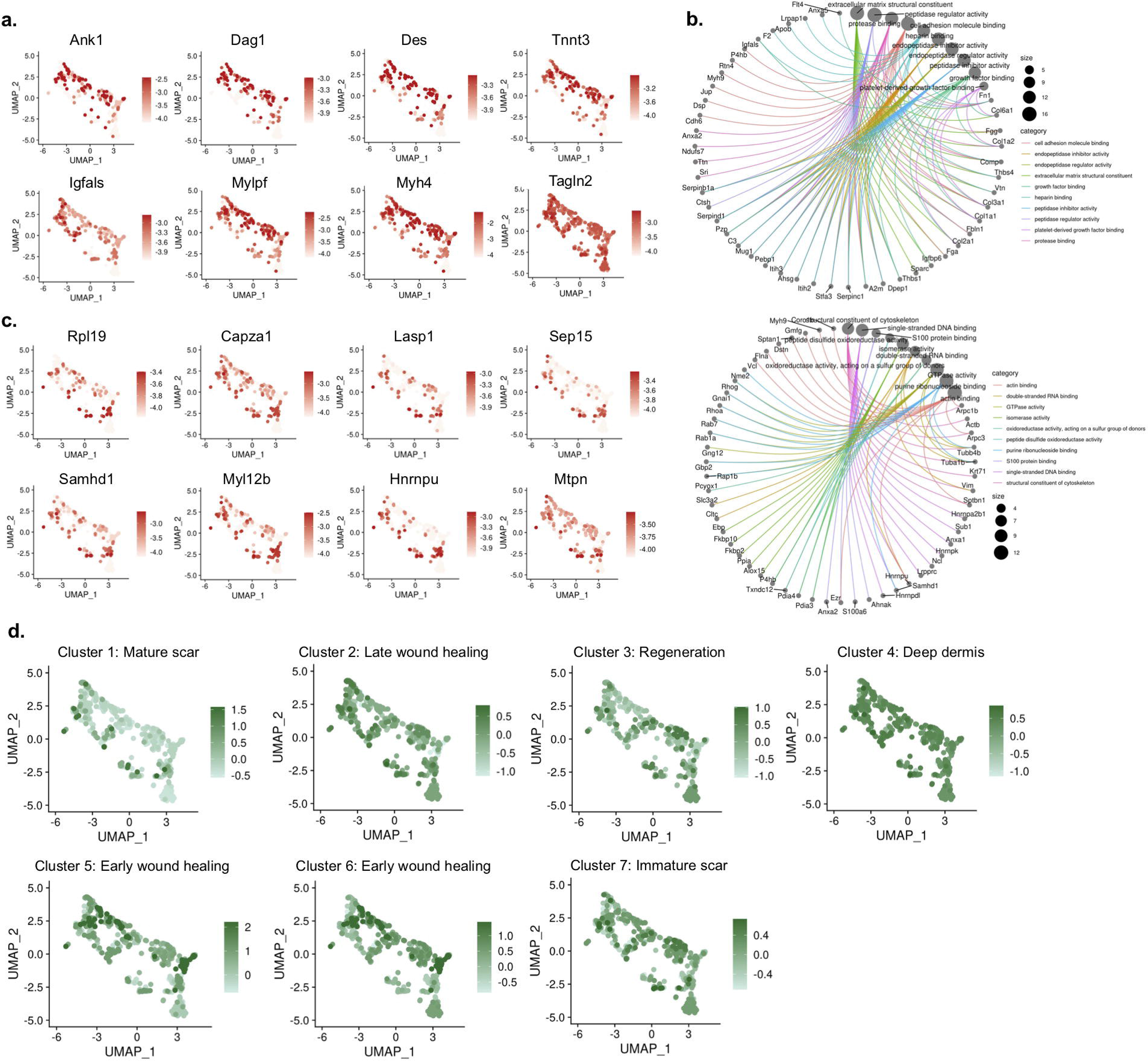
Integration of fibroblast transcriptomic, proteomic, and matrix ultrastructural data. **(A)** Imputation onto scRNA-seq fibroblast manifold of selected proteins with expression positively correlated with pseudotime. **(B)** Functional enrichment analysis of proteins most strongly positively (top) and negatively (bottom) correlated with pseudotime. **(C)** Imputation onto scRNA-seq manifold of proteins with expression most negatively correlated with pseudotime. **(D)** Imputation of average enrichment for each of 7 ECM parameter clusters onto fibroblast scRNA-seq manifold.

We next performed similar imputation analysis of our ECM ultrastructural data using the aforementioned mouse-to-mouse variation in histological parameters (**Figure S4**). Upon initial hierarchical clustering analysis, POD 2 and 7 wounds generally clustered together, suggesting that early wounds exhibited similar provisional matrix features, while verteporfin-treated wounds at POD 30 clustered most closely with unwounded skin (**Figure S4A**), consistent with these wounds regenerating native unwounded skin ECM ultrastructure. Overall, from the 294 total ECM parameters quantified, we identified seven distinct parameter clusters (each comprised of multiple individual parameters) that differentiated the individual samples (**Figure S8A**) and analyzed their enrichment in different wound conditions. Clusters 5 and 6 were most enriched in POD 2 wounds in both treatment groups, suggesting these clusters contained characteristic features of the provisional matrix deposited in early wounds (**Figure S8B,** third row). Consistent with this cluster identity, representative images from these clusters contained largely immature collagen, with a sparse arrangement of short ECM fibers. Using data imputation, we integrated our matrix ultrastructural dataset with the fibroblast pseudotime trajectory to correlate ECM features to specific cell populations. Clusters 5 and 6 were correlated with POD 2 cells in both the scarring and regenerative trajectories (**Figure 3D**), suggesting that provisional matrix deposited in early wounds is similar for both healing outcomes.

While some other clusters were enriched similarly in regenerating and fibrotic wound conditions – for instance, cluster 2, which we termed “late wound healing” features, was most strongly enriched in later (POD 7 +) wounds in both PBS and verteporfin groups (**Figure S8B,** top row right) – several parameter clusters appeared to differentiate fibrotic versus regenerative wounds at different phases. Cluster 1 comprised features most strongly enriched in POD 14 and 30 control (PBS) wounds, suggesting this cluster consisted of ECM features characteristic of mature scars (**Figure S8B,** top row left). Representative images for this cluster had a typical scar-like appearance, with dense, linearly-aligned collagen fibers. Cluster 7, which we termed “immature scar”, had characteristically negative Z-scores in POD 2 wounds and negatively correlated to POD 2 and 7 fibroblasts in the scarring trajectory (**Figure 3D**); this cluster was represented by images of loosely organized, aligned collagen networks (**Figure S8B,** bottom row).

Clusters 3 and 4 were most characteristic of unwounded skin and were also strongly enriched in POD 30 verteporfin-treated wounds. Representative histologic images for cluster 3 contained shorter and broader ECM fibers with a basket-weave/cross-hatched pattern typical of regenerated papillary dermis (**Figure S8B, second row left**). Cluster 4 images, on the other hand, contained aligned, ribbon-shaped fibers consistent with the deep dermis, which were notably absent from POD 30 scars (**Figure S8B, second row right**). While cluster 4 weakly correlated to cells in both arms, with the exception of some POD 14 and 30 scarring trajectory fibroblasts, cluster 3 parameters correlated strongly with fibroblasts in the regenerative trajectory (**Figure 3D**). Ultimately, by leveraging inherent mouse-to-mouse variability and using cell barcoding to perform paired data analysis between our three modalities, we imputed bulk proteomic and ECM ultrastructure data onto trajectories of regeneration and scarring defined at the single-cell level. We were thus able to identify quantitative features of protein expression and tissue architecture that differentiated specific phases of regenerative versus fibrotic tissue repair and correlate these features to transcriptionally distinct fibroblast populations.

Notably, while both proteomic and matrix ultrastructural data exhibited correlation of certain genes or ECM parameters with distinct fibroblast scRNA-seq clusters, such trends were not observed in other cell types. When we performed similar imputation analysis of both bulk data modalities onto myeloid and lymphoid cell scRNA-seq data, there did not appear to be any correlation of the identified fibrotic and regenerative proteomic or ECM features with either myeloid (**Figure S5F and S5G**) or lymphoid cell trajectories (**Figure S6F and S6G**). Instead, proteomic and ECM ultrastructural data imputation supported known roles of these cell populations in wound healing, such as myeloid cell chemotaxis, inflammation, and apoptotic cell clearance (data not shown). This finding is consistent with our initial scRNA-seq pseudotime analysis suggesting that phenotypic differences in fibroblasts, and not in myeloid or lymphoid cells, seem to be the primary driver of the different healing outcomes observed with control versus verteporfin treatment.

A striking difference between PBS- and verteporfin-treated wounds is the emergence of regenerated hair follicles in the latter within 30 days; control wounds typically form bare areas and do not regenerate hair follicles. We wondered if our trajectory-based fibroblast analysis could reveal a master regulator supporting robust regeneration of hair follicles and glands. One gene of particular interest that emerged was *Trps1*, a zinc finger transcription factor whose expression was strongly enriched in the regenerative transcriptional trajectory arm (**Figure 2E**). *Trps1* was previously reported as a novel “master regulator” of Wnt signaling, a pathway well known to play critical roles in hair follicle development, growth, and proliferation.(Fantauzzo and Christiano, 2012; Fantauzzo et al., 2012; Ito et al., 2007) Multiple studies have implicated *Trps1* specifically in hair follicle morphogenesis and aberrant *Trps1* activity has been associated with human genetic conditions characterized by excess hair growth,(Fantauzzo and Christiano, 2012; Fantauzzo et al., 2012; Fantauzzo et al., 2008) making it a promising candidate gene for driving hair follicle regrowth in our regenerating wounds. Importantly, *Trps1* has also been reported to be a key transcriptional regulator of YAP signaling, primarily by competing for shared YAP/TEAD genomic binding sites.(Elster et al., 2018) Given our data suggesting that the regenerative trajectory of wound repair was characterized by decreased mechanotransduction and elevated *Trps1* expression, we postulated that *Trps1* may be an important regulator of wound regeneration.

To examine the dynamics of Trps1 during regenerative and fibrotic wound repair, we first performed immunohistochemical staining to compare expression and localization of Trps1 and YAP in wounds (**Figure 4A**). Nuclear YAP colocalization was comparable between control and verteporfin-treated wounds (**Figure 4B**), consistent with the mechanism of action of verteporfin, which binds and sequesters YAP rather than directly altering its expression. On the other hand, nuclear colocalization of Trps1 was greater in verteporfin-treated than control wounds at both POD 14 and 30 (**Figure 4B**). Furthermore, using AI-based nuclear localization analysis to quantify nuclear localization of Trps1 and YAP in thousands of cells, we found a greater number of high nuclear-Trps1+ fibroblasts in verteporfin-treated wounds (**Figure 4C**). Interestingly, consistent with the proposed role of Trps1 in supporting hair follicle regeneration, clusters of high nuclear Trps1 expression were observed specifically around regenerating hair follicles in POD 30 verteporfin-treated wounds (**Figure 4A**, bottom right panel).

**Figure 4.**
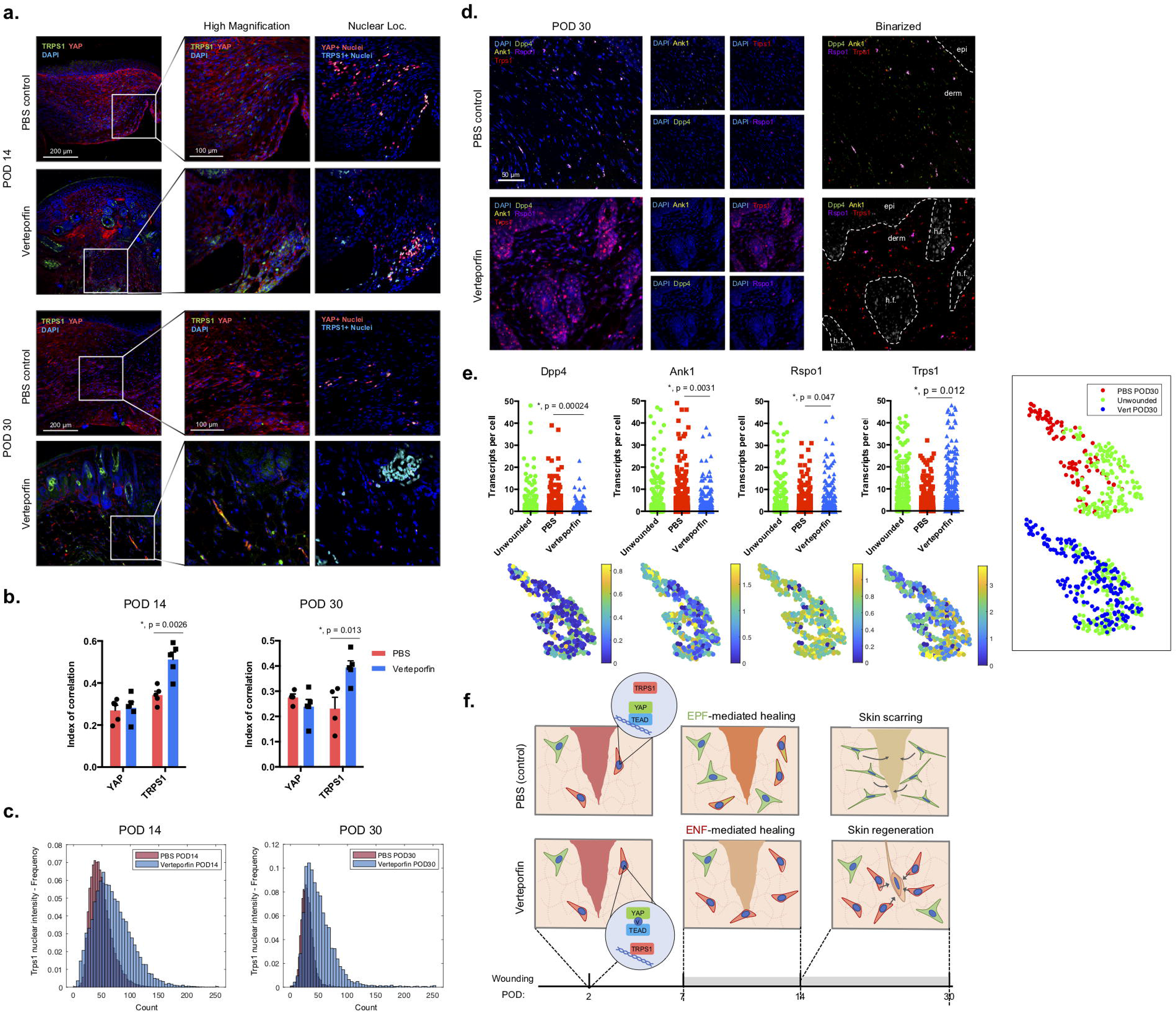
*En-1* lineage-negative fibroblasts promote wound regeneration in the setting of YAP inhibition via increased Trps1 and Wnt signaling. **(A)** Left: Immunofluorescent histology for Trps1 and YAP in control and verteporfin-treated wounds at POD 14 (top) and 30 (bottom). Right: Higher-magnification images of Trps1 and YAP staining (left column) with image processing to highlight nuclear-localized Trps1 (blue) and YAP (red) (right column). **(B,C)** Quantification of Trps1 and YAP nuclear localization by pixel correlation analysis (B) and Biodock AI automated analysis (C). **(D)** Left: RNAscope multiplexed in situ hybridization for Dpp4, Ank1, Rspo1 and Trps1 in control (top) and verteporfin-treated (bottom) wounds. Right: Binarized images highlighting RNA puncta. White dotted line demarcates the quantified regions of interest in the dermis including hair follicles (h.f.). **(E)** Top row panels: Quantification of Dpp4, Ank1, Rspo1, and Trps1 puncta per cell. Bottom row panels: Imputation of quantified RNAscope transcripts per mouse onto corresponding matrix ultrastructural data for that mouse. Right panel: t-SNE embedding of matrix ultrastructural data from unwounded skin and control (top) or verteporfin (bottom) POD 30 wounds. **(F)** Conceptual schematic of scarring (top, YAP- and EPF-mediated) and regenerative (bottom, Trps1- and ENF-mediated) wound repair.

To explore the spatial distribution of *Trps1* activity within wounds at greater depth, we performed multiplexed RNAscope *in situ* hybridization for *Ank1* (a key YAP target gene with enriched protein expression identified in our pro-fibrotic wound trajectory); *Trps1*; *Rspo1* (Wnt agonist identified in our pro-regenerative wound trajectory); and *Dpp4* (CD26), a surrogate marker for the *En-1*-positive fibroblast lineage (unlike *En-1* itself, which is only transiently expressed in EPFs, *Dpp4* is continually expressed following initial *En-1* activation and thus can be used to identify EPFs without requiring genetic lineage tracing (Rinkevich et al., 2015)). Collectively, these genes differentiated the mechanically-activated, pro-fibrotic (*Dpp4*, *Ank1*) and regenerative (*Rspo1*, *Trps1*) pathways elucidated by our scRNA-seq and timsTOF analyses (**Figure 2C and 2E and Figure 3A**). On visual inspection, control (PBS-treated) wound fibroblasts at POD 30 showed greater abundance of *Dpp4* and *Ank1* RNA granules than verteporfin-treated wounds, consistent with a mechanically-activated phenotype. In contrast, verteporfin-treated wounds showed greater abundance of *Rspo1* and *Trps1* RNA granules, particularly around regenerating hair follicles. Quantification of granules across thousands of cells using a custom image analysis pipeline confirmed these trends (**Figure 4E**, top), supporting that ENF-mediated healing (low *Dpp4*) in YAP-inhibited (low *Ank1*) wounds yielded regeneration of functional hair follicles through Wnt-mediated pathway activation (high *Trps1*). Integration of RNAscope with matrix ultrastructural data revealed that *Dpp4* and *Ank1* transcripts were spatially correlated with scar ECM features (**Figure 4E bottom**, first and second columns and right panel, red dots), while *Rspo1* and *Trps1* were correlated with regenerative ECM features (**Figure 4E** third and fourth columns and right panel, green and blue dots).

Collectively, this integrated multimodal analysis supported two trajectories of repair. The first, mediated by EPFs (Dpp4+), is characterized by YAP mechanical activation (high nuclear YAP localization, Ank1+, Trps1-) and results in formation of scar ECM. The second, mediated by ENFs (Dpp4-), is characterized by a lack of YAP mechanical activation (low nuclear YAP localization, Ank1-, Trps1+) and results in regeneration of normal ECM and hair follicles via Wnt pathway activation (**Figure 4F**).

## Discussion

While several studies have examined molecular and cellular characteristics of fibrotic wound repair, none have attempted to delineate divergent mechanisms that define fibrotic versus regenerative pathways of healing. This was recently made possible by our finding that treatment with a small molecule inhibitor of mechanotransduction (verteporfin) prevents postnatal activation of mechanosensitive fibroblasts, yielding true wound regeneration by a pro-regenerative population of fibroblasts (ENFs). We leveraged this phenomenon to directly compare trajectories of fibrotic wound repair (PBS-treated/control wounds) versus regenerative wound healing (verteporfin-treated wounds). By analyzing these two healing outcomes across key timepoints of tissue repair and through multiple molecular lenses (i.e., single-cell transcriptional, translational/proteomic, and matrix ultrastructural), we were able to demonstrate that fibrotic wound repair is characterized by activation of canonical mechanotransduction (YAP) signaling, leading to CD26-positive (i.e., *En-1* lineage-positive) fibroblast-mediated scarring. In contrast, the more regenerative repair observed with verteporfin exhibits disruption of mechanical signaling, with relative enrichment of pathways in CD26-negative (i.e., *En-1* lineage-negative) fibroblasts linked to hair follicle development. Furthermore, we demonstrate that transcriptional divergence between these two trajectories may occur as early as POD 2 in the healing process, with proteomic and histologic divergence following on POD 2/7 and 14, respectively.

These studies provide a framework for leveraging cell surface barcoding and multimodal data imputation to dramatically enhance the granularity and depth of insight that can be gained from bulk datasets. Our multimodal examination also suggested a potential mechanism driving regeneration of dermal appendages and native skin architecture during ENF-mediated repair. Specifically, the transcription factor Trps1, which competes with YAP for binding to shared target chromatin sites, was correlated with a less-fibrotic transcriptional trajectory in the context of YAP inhibition and was spatially linked to *En-1* lineage-negative fibroblasts supporting regeneration of nascent hair follicles. Interestingly, our findings suggest that Trps1 activation and downstream Wnt signaling also occurs in some scarring wound fibroblasts but, in the absence of YAP inhibition, is phenotypically dominated by YAP mechanotransduction signaling. This may indicate that regeneration biologically represents a “default” pathway for repair that is superseded by a tendency towards mechanically-activated fibrosis.

## Acknowledgments

We thank the Stanford Functional Genomics Facility, Stanford Cell Sciences Imaging Facility, and Stanford Shared FACS Facility Cores.

## Author contributions

S.M., H.E.dJ.P, and M.J. conceived, designed, and oversaw the experiments, with suggestions from N.M., R.C., P.K.J., G.G. and M.T.L. S.M., H.E.dJ.P, M.J., K.C., M.F.D., J.D., D.H., M.G., C.A.B., D.S.F., N.M., and R.C. performed wounding experiments, prepared specimens, imaged, and analyzed the data. S.M., H.E.dJ.P., and M.J. wrote the manuscript. P.K.J., D.C.W., G.C.G., and M.T.L. edited the manuscript.;

## Data and materials availability

All data to support the conclusions in this manuscript can be found in the figures. Source data for graphs will be made available in the online version of the paper. Any other data can be requested from the corresponding authors.

## Materials and Methods

### Mice

Mice were housed at the Stanford University Comparative Medicine Pavilion under the care of the Department of Comparative Medicine in the Veterinary Service Center (VSC), in accordance with Stanford APLAC guidelines (APLAC-11048). Ten C57BL/6J mice were acquired from Jackson Laboratory (8 weeks of age) and wounded (see section below) for each experimental time point (POD 0, 2, 7, 14, 30; PBS-or verteporfin-treated).

### Dorsal Excisional Wounding

Experimental mice were wounded in accordance with established protocols.(Galiano et al., 2004) Briefly, mice were anesthetized (2% isoflurane), their dorsal hair was removed with depilatory cream, and the dorsal skin was prepped with alternating betadine and alcohol wipes. Two 6 mm full-thickness circular wounds were placed (extending through the panniculus carnosus) on the dorsum of each animal, roughly 6 mm below the ears and 4 mm lateral to the midline on each side. The wounds were stented open with silicone rings, which were secured around the wound perimeter with adhesive and eight simple interrupted 6-0 nylon sutures (Ethicon) per ring. For mice receiving mechanotransduction inhibitor (verteporfin), 30 μL of verteporfin (1 mg/mL) per wound was injected locally into the wound base; PBS was injected into wounds for vehicle controls. Postoperative analgesia was accomplished with buprenorphine SR 0.05 mg/kg every four hours for three doses, and then as indicated. Dressings were changed every other day under anesthesia. At the indicated post-operative day (POD; 2, 7, 14, or 30), the wounds were harvested by excising a 2 mm ring of tissue around each wound using a 10 mm biopsy punch. One-third of each wound was processed for histology as described below; the remaining two-thirds of each wound were processed for FACS as described below. For unwounded (POD 0) samples, uninjured skin was harvested from the dorsum from a comparable region to wound samples and split between histology and FACS as for wounds.

### Fluorescence Activated Cell Sorting

The dorsal skin was harvested using dissecting scissors by separation along fascial planes. Next, the subcutaneous fat was trimmed with a scalpel, and the skin was rinsed in betadine, followed by 5 rinses in cold PBS. To achieve a cell suspension, the harvested skin was finely minced using sharp scissors, enzymatically digested (Liberase DL, 0.5 mg/mL, 1 hour), and filtered through a 40 μm nylon mesh. Cells were isolated from experimental mice via a previously reported FACS strategy. Briefly, a lineage gate (Lin) for hematopoietic (CD45, Ter-119), endothelial (CD31, Tie2), and epithelial (CD326, CD324) cell markers was used as a negative gate to isolate fibroblasts (Lin^−^) and non-fibroblasts (Lin^+^).

### Single-cell Transcriptomic Analysis

Cell suspensions were labeled with TotalSeq Series B hashtag oligonucleotide-labeled antibodies (BioLegend). Single-cell RNA-seq (scRNA-seq) was then performed at the Stanford Functional Genomics Facility (SFGF) for droplet-based microfluidics using the 10x Chromium Single Cell platform (Single Cell 3’ v3, 10x Genomics, USA). Droplets of the cellular suspensions, reverse transcription master mix, and partitioning oil were mixed, loaded onto a single cell chip, and processed on the Chromium Controller. Reverse transcription was performed, and cDNA was amplified using a BioRad C1000 Touch thermocycler, with cDNA size selected using SpriSelect beads (Beckman Coulter, USA). An Agilent Bioanalyzer High Sensitivity DNA chip was used to analyze cDNA for qualitative control purposes; cDNA was then fragmented using the proprietary fragmentation enzyme blend for 5min at 32°C, followed by end repair and A-tailing at 65°C for 30 min. DNA was double-sided size selected using SpriSelect beats. Sequencing adaptors were ligated to the cDNA at 20°C for 15min. cDNA was amplified using a sample-specific index oligo as primer, followed by another round of double-sided size selection using SpriSelect beads. Final libraries were analyzed on an Agilent Bioanalyzer High Sensitivity DNA chip for qualitative control purposes. Libraries were sequenced on a HiSeq 4000 Illumina platform targeting 50,000 reads per cell.

Base calls were converted to reads using the Cell Ranger (10X Genomics; version 3.1) implementation mkfastq and then aligned against the Cell Ranger mm10 reference genome, available at: http://cf.10xgenomics.com/supp/cell-exp/, using Cell Ranger’s count function with SC3Pv3 chemistry and 5,000 expected cells per sample. Cell barcodes representative of quality cells were differentiated from apoptotic cell barcodes or background RNA based on a threshold of having at least 200 unique transcripts profiled, less than 100,000 total transcripts, and less than 10% of their transcriptome of mitochondrial origin. Unique molecular identifiers (UMIs) from each cell barcode were retained for all downstream analysis, normalized with a scale factor of 10,000 UMIs per cell, and subsequently natural log transformed with a pseudocount of 1 using the R package Seurat (version 3.1.1).(Chen et al., 2013) The first 15 principal components of the aggregated data were then used for uniform manifold approximation and projection (UMAP) analysis.(Gulati et al., 2020) Cell annotations were ascribed using SingleR (version 3.11) against the Mouse-RNAseq reference dataset, available at https://rdrr.io/github/dviraran/SingleR/man/mouse.rnaseq.html.

Cell-type marker lists were generated with two separate approaches: first, we employed Seurat’s native *FindMarkers* function with a log fold change threshold of 0.25 using the ROC test to assign predictive power to each gene; second, we employed a characteristic direction analysis to better account for the mutual information contained within highly correlated predictive genes.(Chen et al., 2013)

### Pseudotime Analysis

Pseudotime analysis was performed using the Monocle 3 package in R (version 3 0.2.0).(Trapnell et al., 2014) Counts for individual cells were preprocessed using PCA with 15 dimensions following log-normalization. Dimensional reduction was performed using a UMAP reduction with min_dist = 0.5, n_neighbors = 30, and repulsion.strength = 2.0. Cells were then clustered using Monocle 3’s Louvain implementation with a resolution of 1e-5. A principal graph was then learned from the reduced dimension space using reversed graph embedding with default parameters, and cell order selection was made from the two elements at either end of the trajectory. Pseudotime trajectory heatmaps were created using the Monocle 2 package in R.

For fibroblast, myeloid, and lymphoid cell subseurat’s, we identified a root starting point to calculate pseudotime values for each cell within the dataset. Then, we then used the R function *cor* to correlate pseudotime values with gene expression levels, protein expression z scores, and matrix histology levels. These correlation values were used to determine the genes, proteins, and matrix parameters that either increased (positive correlation) or decreased (negative correlation) with pseudotime. Correlation values in the top 1% of positively or negatively correlated genes were chosen for downstream analyses. Integration of timsTOF proteomic and ECM ultrastructural data onto pseudotime scRNA-seq manifolds was performed pairwise between modalities on a per-mouse basis without replacement for missing values.

### CytoTRACE Analysis

We utilized the recently developed bioinformatics tool CytoTRACE to compare differentiation states among cells in our dataset (https://cytotrace.stanford.edu/).(Gulati et al., 2020) This tool analyzes the number of uniquely expressed genes per cell, as well as other factors like distribution of mRNA content, to calculate a score assessing the differentiation and developmental potential of cells. This analysis was performed using default parameters for each cell in our dataset.

### RNA Velocity Analysis

RNA velocity analysis was performed using the scVelo package.(Bergen et al., 2020) scVelo uses a likelihood-based dynamical model to solve the full transcriptional dynamics of spliced and unspliced mRNA kinetics of each gene. RNA velocity analysis allowed us to identify transient cellular states in our dataset and to predict the directional progression of transcriptomic signatures along the identified trajectories. These predictions are based on gene-specific rates of transcription, splicing, and degradation of mRNA to estimate each cell’s position in their own underlying differentiation process. The RNA velocity across all genes is then projected as a stream of arrows on the UMAP embedding.

### timsTOF Shotgun Proteomic Analysis

Cells for proteomic analysis were pelleted using a swinging-bucket rotor centrifuge (500 x g, 5 minutes, 4 ºC). Supernatant was removed, and pellets were snap frozen on dry ice, then stored at –80 ºC until protein purification. Peptides were purified from pellets using the PreOmics iST kit (Martinsried, Germany) following the kit protocol. Samples were homogenized by passing through QIAshredder spin columns (QIAGEN) following lysis. Peptide concentration was quantified with a plate reader using the Pierce Quantitative Colorimetric Peptide Assay (Thermo Scientific). Finally, peptide samples were diluted to normalize concentration and 50 μg of each was loaded into the timsTOF analyzer. The timsTOF Pro (Bruker) was operated in PASEF mode using Compass Hystar 5.0.36.0. All raw files were analyzed by MaxQuant v1.6.6 software using the Andromeda search engine. Functional enrichment and network analyses were performed on positively and negatively correlated pseudotime genes using EnrichR.

### Histology and Immunofluorescence

Samples were placed into histology cassettes and fixed by incubation in 5% buffered formalin phosphate for 16 hours at 4 ºC. The samples were washed with PBS, dehydrated through serial ethanol washes, cleared with xylene, infiltrated with paraffin through serial incubations, and embedded in paraffin. Sections were cut at a thickness of 8 μm and incubated at 37 ºC overnight to affix sections to slides prior to staining. Picrosirius Red and Masson’s Trichrome staining was performed per standard protocols from the manufacturer (Abcam).

For immunofluorescent staining, samples were cleared with xylene, re-hydrated, and treated with antigen retrieval buffer per established protocols (Abcam ab970). Samples were then permeabilized in 0.25% Triton-X (15 minutes), blocked for 2 hrs (Powerblock), and stained overnight with primary antibodies in 0.1x Powerblock (YAP Santa Cruz Biotechnology sc-101199, Trps1 Proteintech 20003-1-AP). Samples were then washed 3 times with 0.1x Powerblock, stained for 1 hr with secondary antibodies (Invitrogen), washed another 3 times, and finally mounted with Fluoromount-G containing DAPI. Slides were imaged using a Leica TCS SP5 confocal microscope. Nuclear localization analysis of YAP and Trps1 was performed using the EzColocalization Fiji and BioDock AI Nuclear Segmentation modules (www.biodock.ai).(Stauffer et al., 2018)

### Quantitative Analysis of Connective Tissue Ultrastructure

Picrosirius Red-stained histologic sections derived from unwounded skin and wounds (5 PBS- and 5 verteporfin-treated, 2 wounds per mouse, 3 sections per slide) were imaged in tiles at 40x magnification. Next, images were normalized using the RGB histogram method (Stain Normalization Toolbox, University of Warwick) and color deconvoluted using the algorithm previously described by Ruifrok et al.,(Ruifrok and Johnston, 2001; Ruifrok et al., 2003) wherein each pure stain is characterized by absorbances within three RGB channels. Ortho-normal transformation of the histology images produced individual images corresponding to each color’s individual contribution to the image. Applied to birefringent Picrosirius Red images (green to red color under polarized light depending on packing of fiber bundles), this technique produced deconvoluted red and green images corresponding to mature and immature connective tissue fibers, which were then analyzed independently. Noise reduction of deconvoluted fibers was achieved using an adaptive Wiener filter in Matlab 2019a (wiener2 function), which tailors itself to the local image variance within a pre-specified neighborhood (3-by-3 pixels in our application). The filter preferentially smooths regions with low variance, thereby preserving sharp edges of fibers. Smooth images were then binarized using the im2bw command and processed through erosion filters with diamond-shaped structuring elements to select for fiber-shaped objects. Finally, the fiber network was “skeletonized” using the bwmorph command and various parameters of the digitized map (fiber length, width, persistence, alignment, etc.) were measured using the regionprops command. Dimensionality reduction of quantified fiber network properties by t-distributed stochastic neighbor embedding (t-SNE) was achieved using the default tsne (distance metric specified as Euclidian distance) command in Matlab. Hierarchical clustering was performed by first filtering low variance parameters (bottom 25%) and then calculating Euclidean distances. Matlab scripts containing our fiber quantification pipeline are available at the following Github repository: https://github.com/shamikmascharak/Mascharak-et-al-ENF.

### Spatial Transcriptomic Analysis

RNAscope was used to evaluate spatial distributions of mechanically-activated fibrotic (Ank1+, CD26+) and non-activated regenerative (Trps1+, Rspo1+) fibroblasts. Paraffin-embedded slides were processed in house by the manufacturer (ACD, Newark CA) for multiplexed *in-situ* hybridization. Slides were then imaged on a Leica TCS SP8 confocal microscope at 40x magnification. Spatial transcriptomic patterns were analyzed using a custom Matlab pipeline. Briefly, confocal images were split into separate fluorescent channels and images of nuclei (DAPI) and RNA puncta (Dpp4 Opal 520, Ank1 Opal 570, Rspo1 Opal 620, Trps1 Opal 690) were binarized using shaped-based filtration. Next, regions of interest were traced around nuclei with an additional 10-pixel berth in order to capture somatic transcripts. Finally, RNA puncta per cell were quantified across all cells and images for each mouse. Matlab scripts containing our RNAscope analysis pipeline are available at the following Github repository: https://github.com/shamikmascharak/Mascharak-et-al-ENF.

