## Supplementary Materials for "Divergent molecular signatures of regeneration and fibrosis during wound repair"

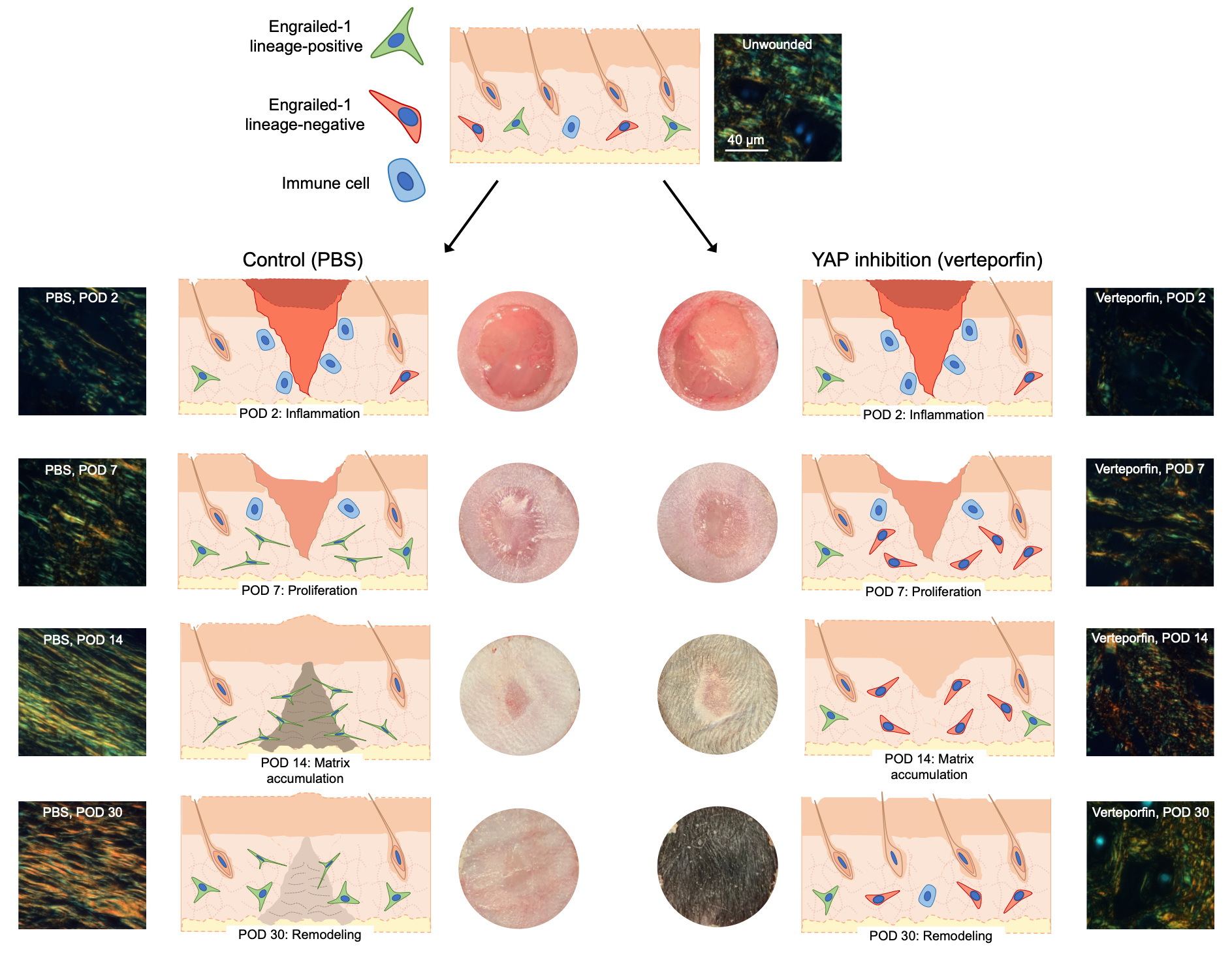

Figure S1: Scarring wound repair versus wound regeneration in the setting of YAP inhibition. Schematics (middle panels), gross photos (inner columns), and Picrosirius Red histology (outer columns) of PBS- (control; left) and verteporfin-treated (right) wounds. YAP inhibition with verteporfin yields wound regeneration by POD 30, with full recovery of hair follicles and glands.

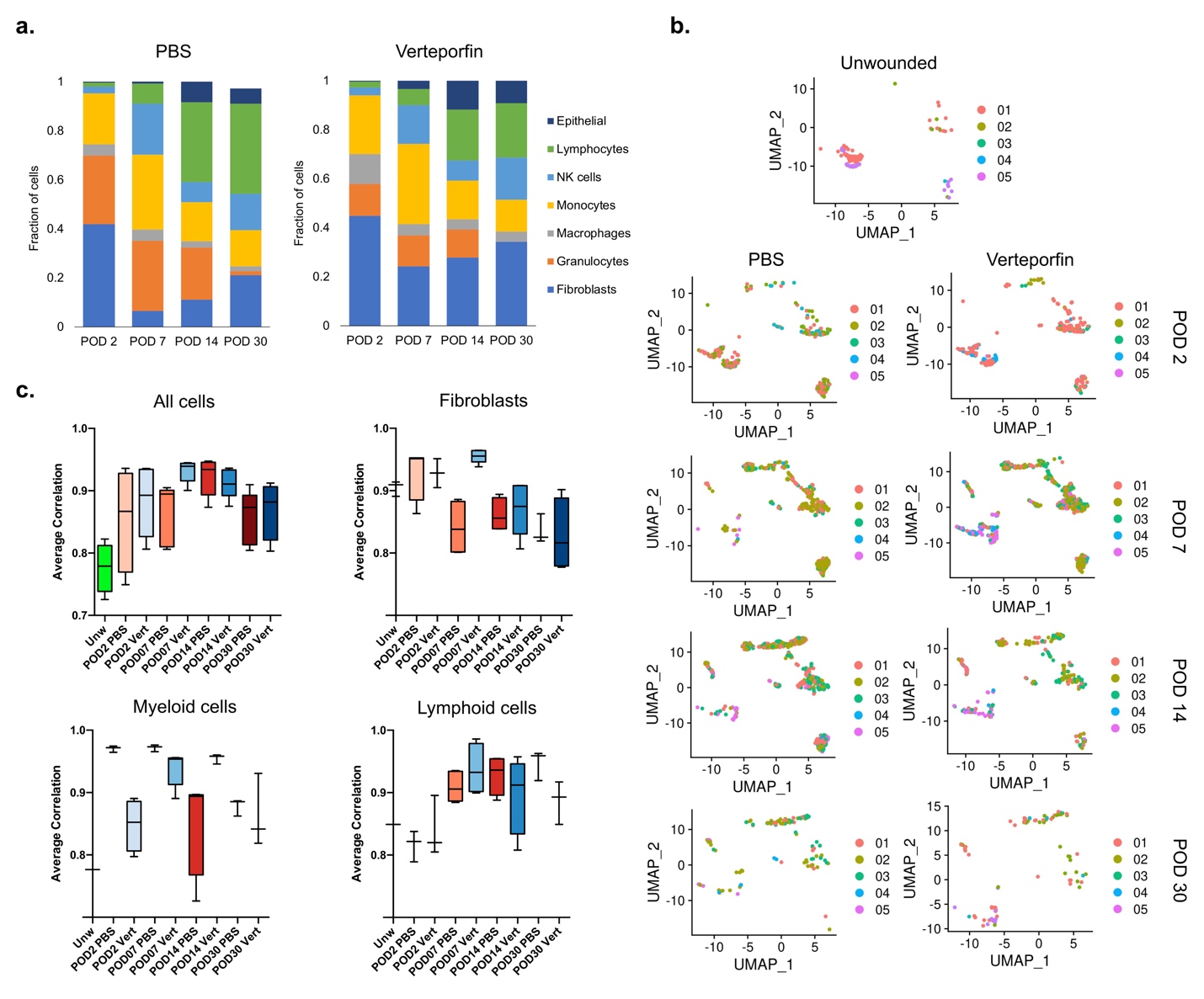

**Figure S2: Single cell RNA sequencing-based comparison of scarring and regenerating skin wounds. (A)** Quantification of cell type abundances per POD in control (left) and verteporfin-treated (right) wounds, inferred from scRNA-seq. **(B)** Fibroblast UMAP showing cells from unwounded skin (top panel) and control (left column) or verteporfin (right column) wounds at POD 2 (second row), 7 (third row), 14 (fourth row), and 30 (bottom row), each colored by individual mice (*n* = 5 mice per timepoint/condition). **(C)** Mouse-to-mouse correlation of scRNAseq data for all cells (top left) or fibroblasts (top right), myeloid cells (bottom left), or lymphoid cells (bottom right) only for all timepoints/conditions.

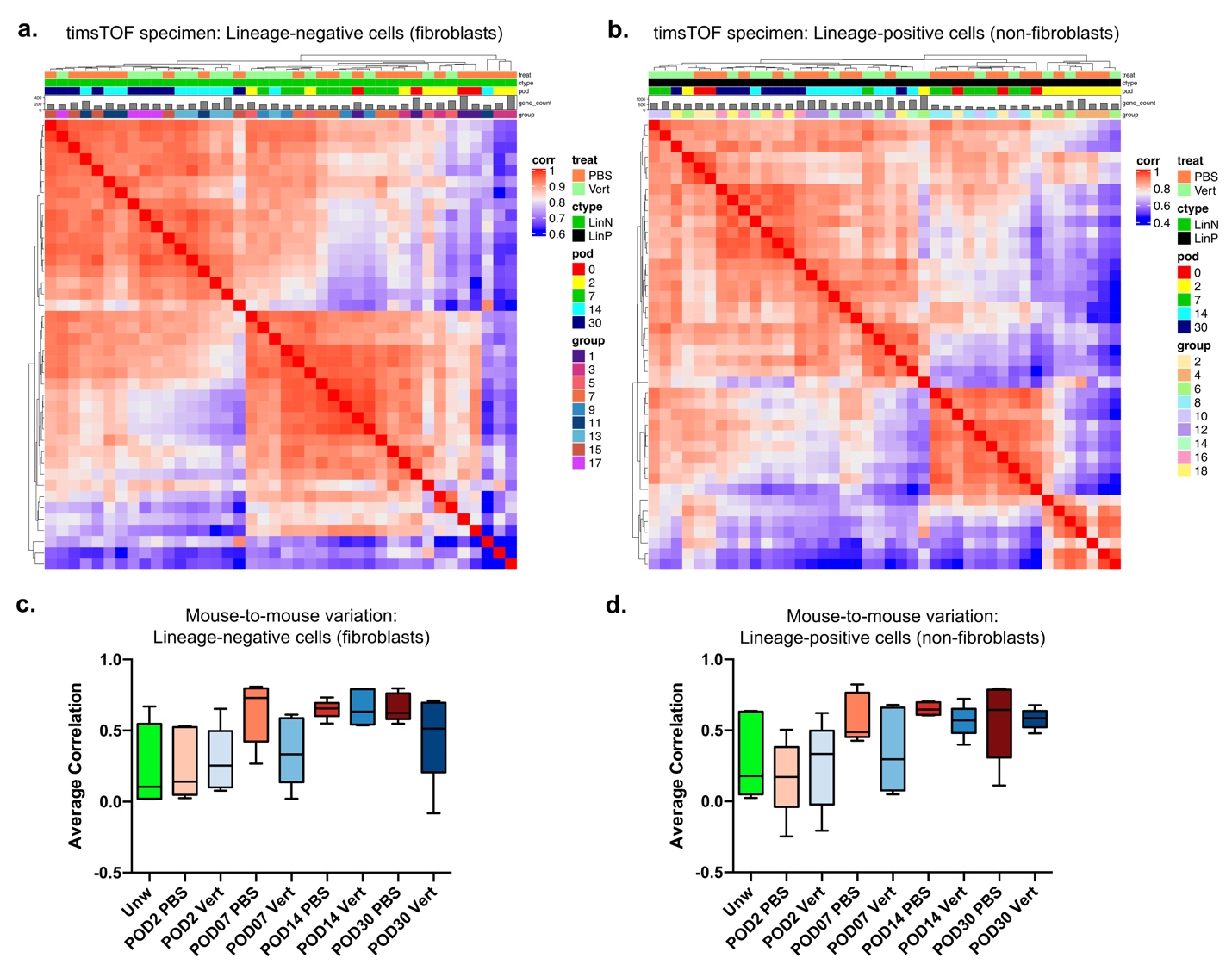

**Figure S3: Mouse-to-mouse heterogeneity of timsTOF proteomic profiles. (A,B)** Correlation heatmaps of bulk timsTOF data derived from lineage-negative (fibroblasts, A) and lineage-positive (non-fibroblasts, B) FACS-isolated cells at POD 0 (unwounded), 2, 7, 14, and 30. Each column represents protein enrichment averages for a single mouse. **(C,D)** Mouse-to-mouse correlation of timsTOF data for lineage-negative (C) and lineage-positive (D) cells.

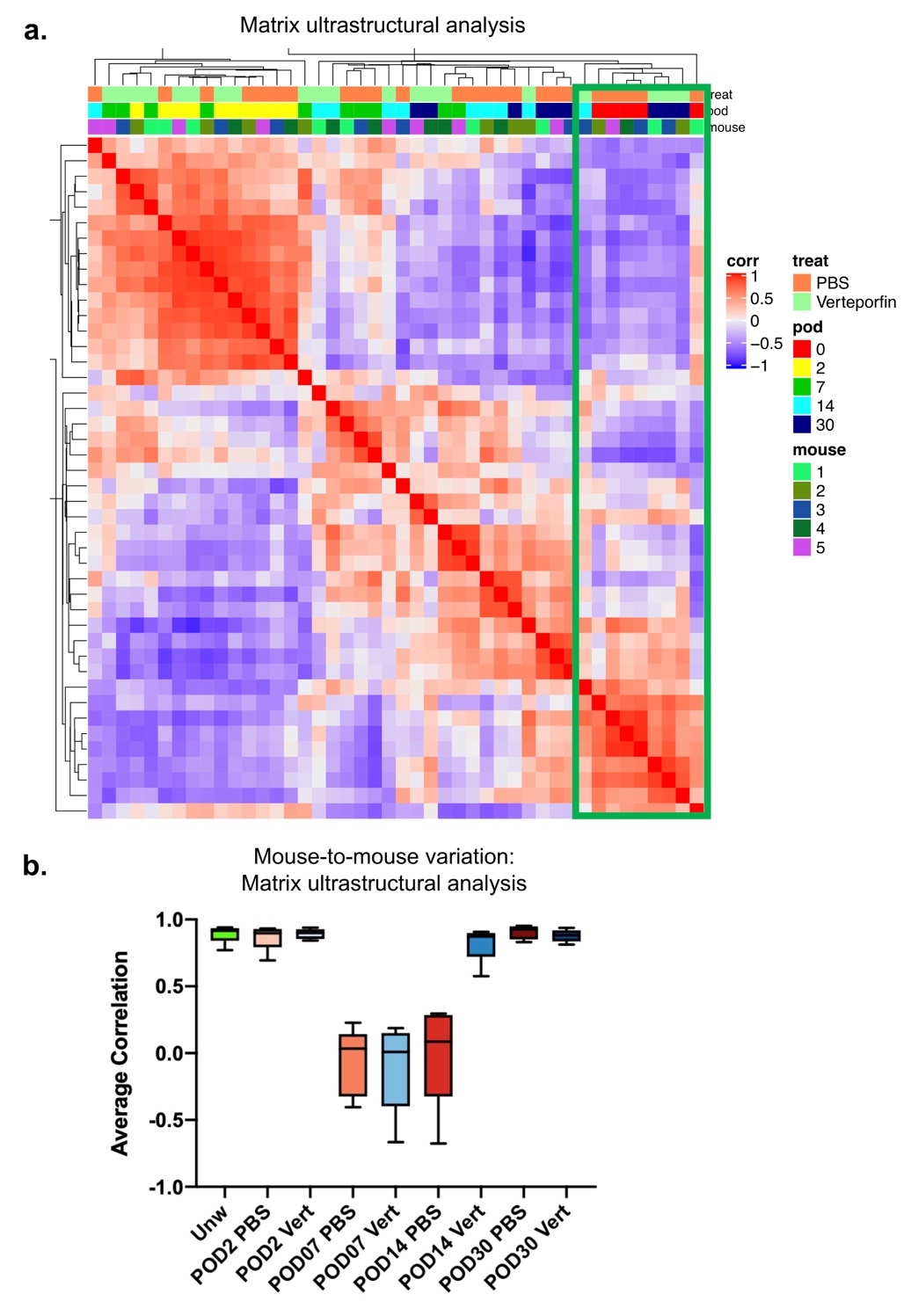

**Figure S4: Mouse-to-mouse heterogeneity of ECM ultrastructural data. (A)** Correlation heatmap of quantified ECM data for control- and verteporfin-treated wounds at POD 0 (unwounded), 2, 7, 14, and 30. Each column represents matrix parameter averages for a single mouse. Green box highlights several verteporfin-treated wounds at POD 30 clustered together with unwounded skin. **(B)** Mouse-to-mouse correlation of matrix ultrastructural data.

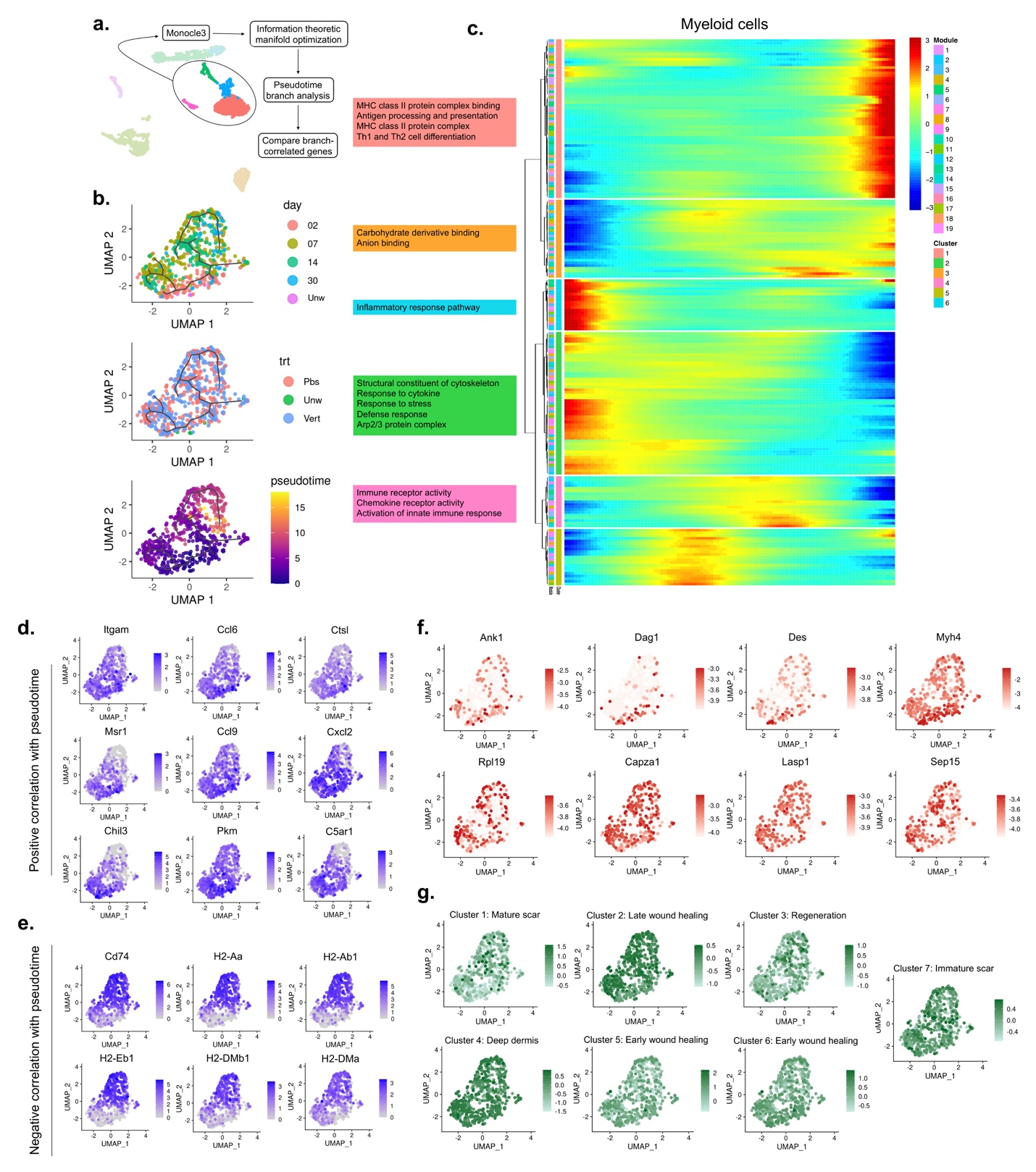

**Figure S5: Multimodal analysis of myeloid cells during skin scarring and regeneration. (A)** Schematic outlining manifold generation, optimization, and branchpoint analysis for myeloid cells (macrophages, monocytes, granulocytes, etc.) in Monocle3. **(B)** Myeloid cell manifold colored by POD (top), treatment group (middle), and pseudotime value (bottom), showing no diverging behavior in scarring (PBS control) and regenerating (verteporfin) wounds. **(C)** Clustering of myeloid cells by pseudotemporal expression pattern. Colored boxes (left) contain functional enrichment terms for the corresponding gene cluster. **(D,E)** mRNA expression of selected genes negatively (D) and positively (E) correlated with pseudotime, demonstrating similar activation of early inflammation (D) and later antigen presentation-related (E) pathways in scarring and regeneration. **(F,G)** Imputation of matrix ultrastructural (F) and proteomic (G) data onto the manifold in (B) did not differentiate a scarring and a pro-regenerative signature, suggesting that myeloid cell function was largely unchanged in the context of verteporfin treatment.

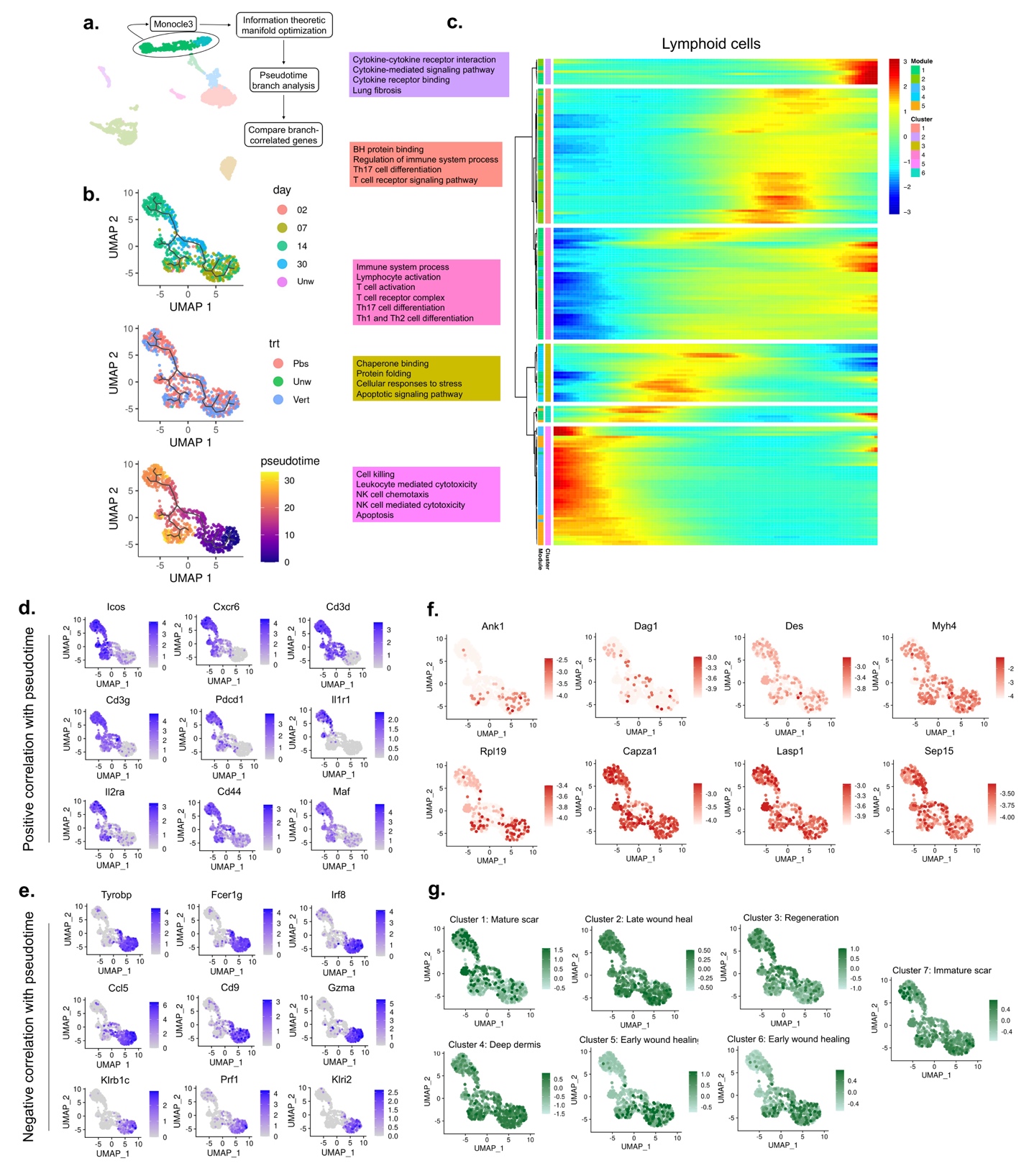

**Figure S6: Multi-modal analysis of lymphoid cells during skin scarring and regeneration. (A)** Schematic outlining manifold generation, optimization, and branchpoint analysis for lymphoid cells (B/T lymphocytes, NK cells) in Monocle3. **(B)** Lymphoid cell manifold colored by POD (top), treatment group (middle), and pseudotime value (bottom), showing no diverging behavior in scarring (PBS control) and regenerating (verteporfin) wounds. **(C)** Clustering of lymphoid cells by pseudotemporal expression pattern. Colored boxes (left) contain functional enrichment terms for the corresponding gene cluster. **(D,E)** mRNA expression of selected genes positively (D) and negatively (E) correlated with pseudotime, demonstrating similar activity of early NK cells (E) and later B/T lymphocytes (D) in scarring and regeneration. **(F,G)** Imputation of matrix ultrastructural (F) and proteomic (G) data onto the manifold in (B) did not differentiate a scarring and a pro-regenerative signature, suggesting that lymphoid cell function was largely unchanged in the context of verteporfin treatment.

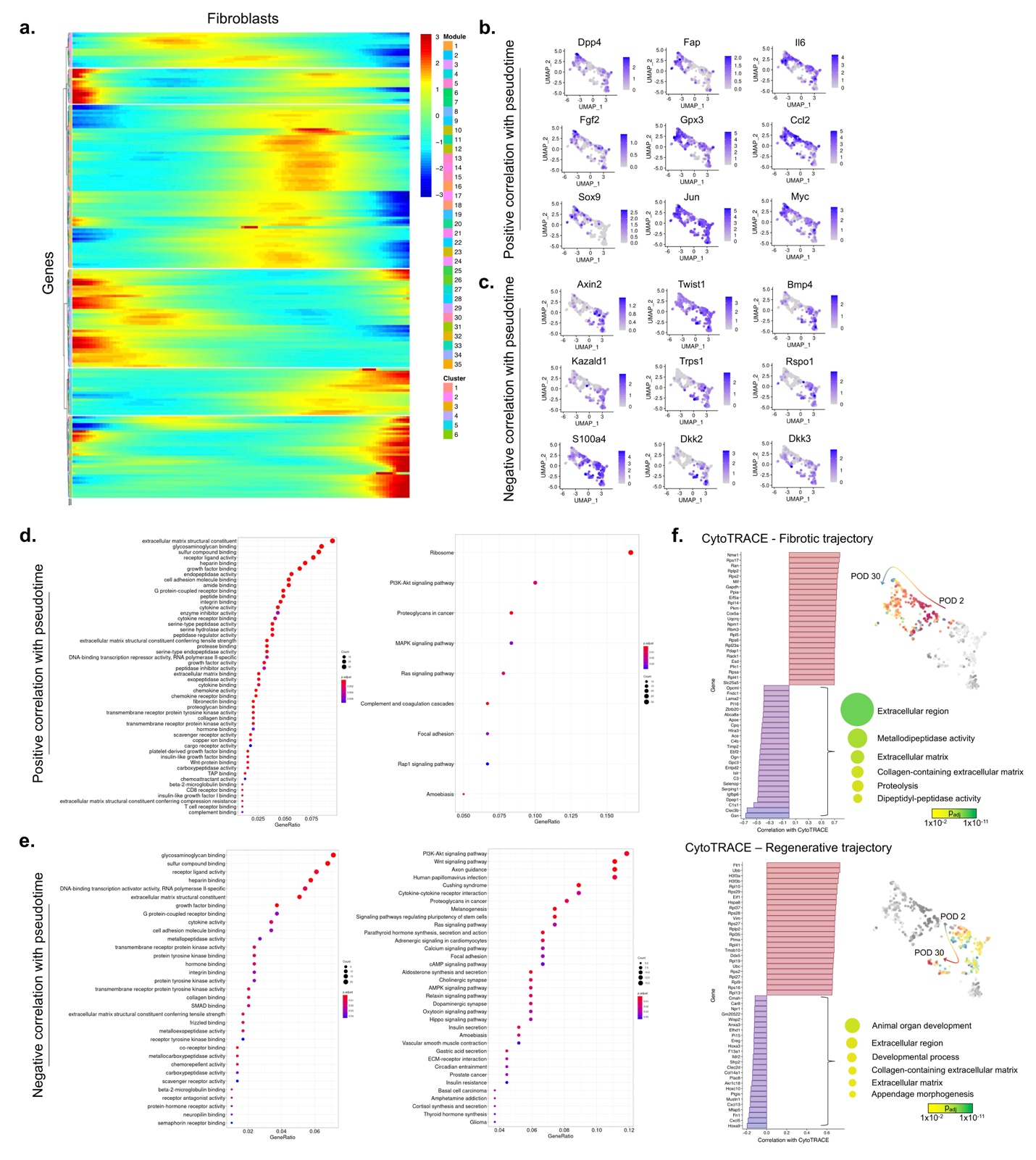

**Figure S7: Pseudotime correlation and CytoTRACE analysis of fibroblasts in skin scarring and regeneration. (A)** Clustering of fibroblasts by pseudotemporal expression pattern. **(B)** mRNA expression feature plots for selected genes positively correlated with pseudotime. **(C)** EnrichR Gene Ontology analysis of top 1% of genes positively correlated with pseudotime revealed signatures consistent with mechanical activation/fibrosis. **(D)** mRNA expression feature plots for selected genes negatively correlated with pseudotime. **(E)** Functional enrichment analysis of the top 1% of genes negatively correlated with pseudotime revealed signatures consistent with stem cell and development pathways (e.g., Wnt, BMP signaling). **(F)** Waterfall plots showing genes positively (red) and negatively (blue) correlated with CytoTRACE score for cells in the fibrotic (top) and regenerative (bottom) fibroblast trajectories. Right panels show functional enrichment results for genes correlated with decreasing CytoTRACE score (i.e., with fibroblast differentiation).

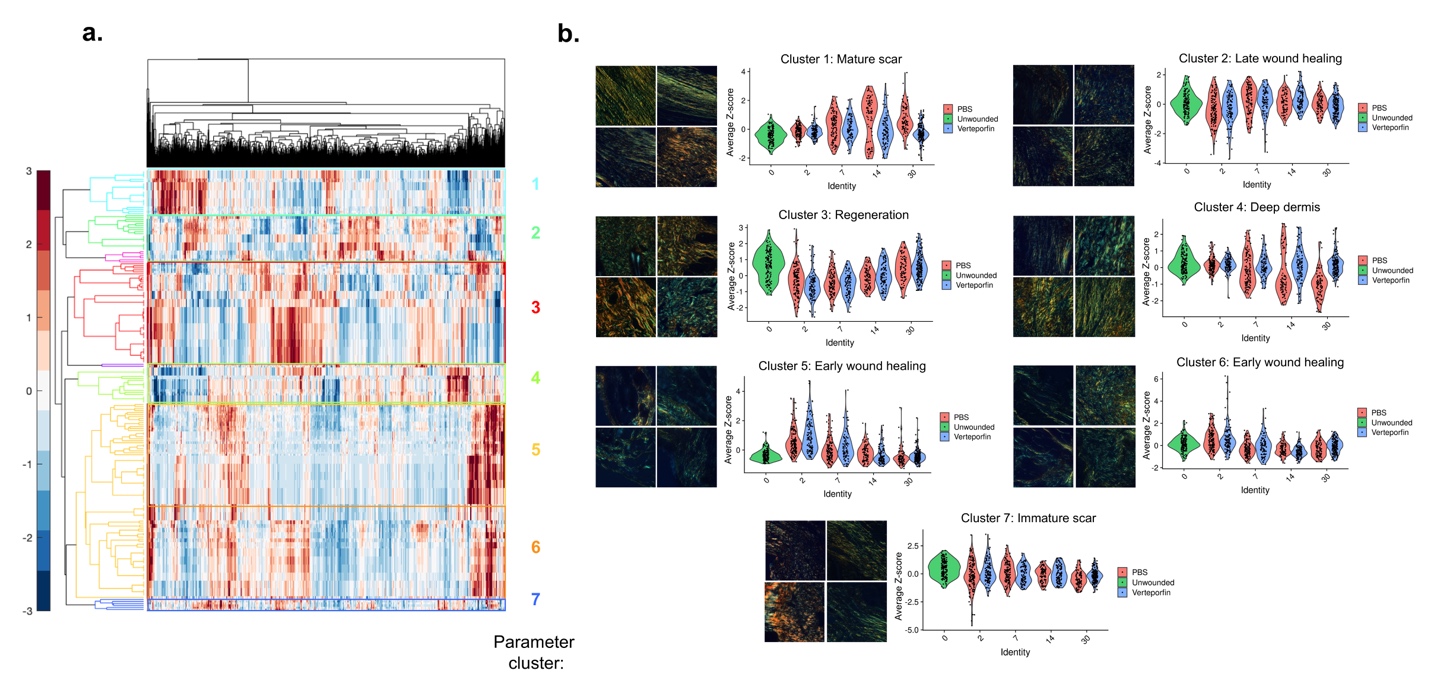

**Figure S8: Hierarchical clustering of matrix ultrastructural parameters. (A)** Clustergram of over 1000 matrix histology images (columns) differentiated on the basis of 7 major cluster parameters (colored row brackets). **(B)** Quantification of average parameter cluster values per timepoint/condition (right panels) with corresponding representative histology images (left panels) for each of the 7 major parameter clusters.

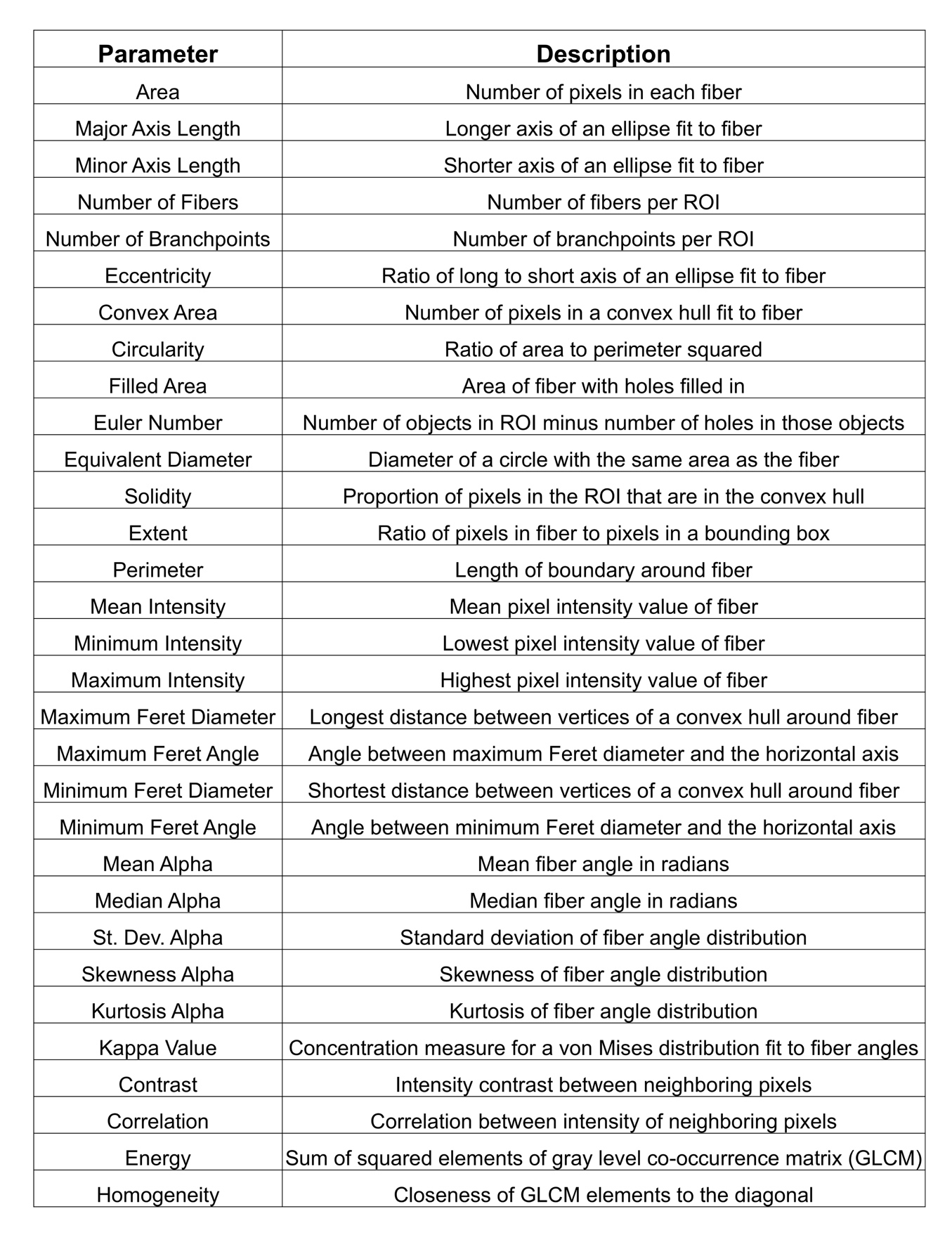

**Table S1: Descriptions of quantified matrix parameters.** Note that each parameter was quantified for color deconvoluted mature (red) and immature (green) Picrosirius Red-labeled fibers. For parameters quantified across all fibers in an image (thus producing a distribution), the mean and standard deviation were both used.
